# Integrating AI and molecular modeling for structural prediction of a closed state of the hERG channel

**DOI:** 10.64898/2026.04.20.719540

**Authors:** Catherine Upex, Thomas Osborne, Giovanni Biglino, Jules Hancox, Robin A Corey

**Affiliations:** Bristol Medical School (THS), Level 7, Bristol Royal Infirmary, Upper Maudlin Street, Bristol, BS2 8HW; Bristol Medical School (THS), Cardiovascular Research Laboratories, Biomedical Sciences Building, University Walk, Bristol, BS2 8HW; School of Psychology and Neuroscience, Biomedical Sciences Building, University Walk, Bristol, BS8 1TD

## Abstract

The voltage-gated potassium channel hERG (Kv11.1) plays a central role in cardiac repolarisation by mediating the rapid delayed rectifier K^+^ current (I_Kr_). Blockage of hERG by small molecules can lead to delayed repolarisation, QT interval prolongation, and potentially fatal arrhythmias, making the channel a critical focus in drug safety screening. Despite extensive pharmacological and electrophysiological characterisation, a complete structural understanding of hERG gating remains limited by the absence of an experimentally determined closed-state structure.

Here, we use AI-based structural modelling to predict and compare candidate closed conformations of hERG. Building on recent work in which AlphaFold2 (AF2) predictions guided by engineered structural templates captured closed and inactivated states, we applied the emerging protein structure predictor, Chai-1, which employs a single-sequence, language model-based approach independent of multiple-sequence alignments. The resulting Chai-1 hERG model was compared with the AF2-derived closed structure, a homology model based on the *Rattus norvegicus* EAG channel, and an experimentally resolved open-state cryo-EM structure. We assessed these models using a combination of all-atom and coarse-grained molecular dynamics simulations, analysing protein dynamics, pore geometry, gating residue orientation, hydration, and lipid interactions. The Chai-1 and AF2 models displayed strong structural and dynamic agreement, both adopting compact, non-conductive conformations consistent with a physiologically closed state. Our data reveal insights into VSD dynamics, as well as suggesting a state dependence for ceramide binding at the previously identified M651 residue.

Our findings support the validity of AI-derived closed-state hERG models and underscore the growing potential of deep learning-based protein structure prediction to identify previously uncharacterised, pharmacologically relevant conformations of membrane proteins. Further, our Chai-1 derived closed state model expands our structural insights into hERG gating and may have utility for investigation of drug-hERG interactions.

## Introduction

The voltage-sensitive potassium (K^+^) channel Kv11.1, also known as hERG1 (hereafter hERG), plays a critical role in the repolarisation phase of the cardiac action potential by mediating the rapid component of the delayed rectifier K^+^ current (I_Kr_) (*1*). Owing to this essential physiological role, perturbations to hERG function can have serious clinical consequences. hERG is notably promiscuous in its pharmacology; blockade of the channel by small molecules leads to delayed repolarisation of the cardiac action potential, manifesting as QT interval prolongation on an electrocardiogram – a condition referred to as drug-induced long QT syndrome (diLQTS) – which can precipitate fatal cardiac arrhythmias (*2*–*4*). Consequently, understanding how novel compounds interact with hERG is a critical aspect of early drug discovery and safety evaluation.

Despite extensive interest in hERG as a pharmacological target and anti-target, structural information on the channel across its full conformational landscape (open, closed, and inactivated states) remains incomplete. Several open-state structures have been resolved using cryo-electron microscopy (cryo-EM), including complexes with high-affinity blockers (*5*) (*6*) (*7*). In addition, a non-conductive state – interpreted as inactivated – has recently been elucidated by exploiting the channel’s K^+^-sensitive inactivation mechanism (*6*). However, an experimentally derived closed-state structure remains elusive, with current best practice using homology modelling based on the closely related *Rattus norvegicus* EAG channel (PDB: 5K7L(*8*)) (*9*), or derived using a molecular dynamics (MD) relaxed open structure (*10*). The absence of this key conformation limits our understanding of gating transitions and constrains efforts to model state-dependent drug interactions, limiting our ability to screen drugs for hERG blockage.

The hERG channel exhibits a canonical voltage-gated potassium channel architecture (*6*). It assembles as a tetramer in which each subunit contributes six transmembrane helices (S1-S6). The S1-S4 helices form the voltage-sensing domain (VSD) (**Figure 1:** yellow), which detects changes in membrane potential and drives conformational rearrangements. These movements are transmitted through the intracellular C-linker (**Figure 1**: cyan), that couples the VSD to the pore domain (**Figure 1**: blue). The pore domain is formed by the S5 and S6 helices from each subunit, creating the central ion conduction pathway. Within this region, the S6 helix (**Figure 1**: green) lines the inner pore and plays a key role in gating, while the selectivity filter (**Figure 1**: pink), located at the extracellular end of the pore, confers potassium ion selectivity. Together, these structural elements coordinate to underpin the distinctive gating kinetics and ion conduction properties of hERG.

**Figure 1:**
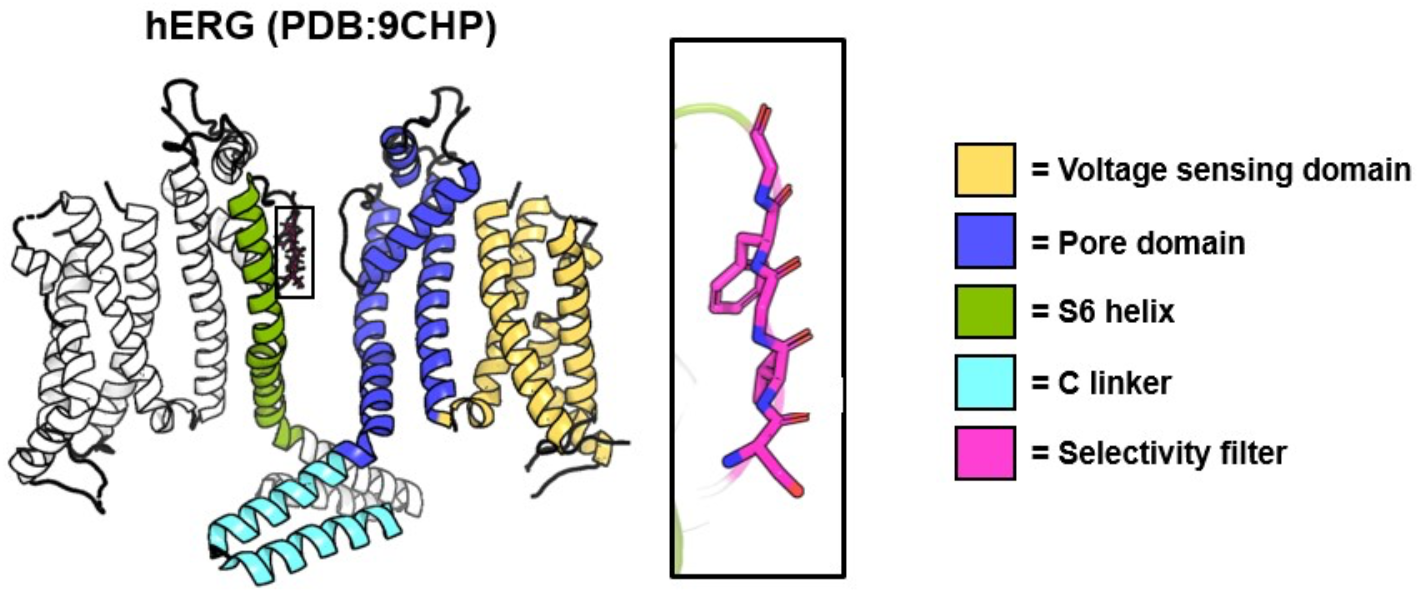
Structural architecture of the hERG channel. Shown are two subunits from the cryo-EM structure of hERG determined in 300 mM K^+^ (PDB: 9CHP). Key regions are highlighted, including the voltage-sensing domain (yellow), C-linker (cyan), and pore domain (blue). The pore domain contains the S6 helix (green) and the selectivity filter (pink).

Advances in computational protein structure prediction have provided new avenues for addressing the gaps in our current structural understanding of the hERG channel. AlphaFold2 (AF2), a deep learning model that integrates multiple-sequence alignments (MSAs) to predict 3D protein structures, has been transformative in structural biology, often achieving near-experimental accuracy even in the absence of close homologues(*11*). AF2 and other AI-based modelling methods have been applied to look at a range of protein targets, including a range of voltage-gated ion channels (*12, 13*). Recent work using AF2 with engineered structural templates successfully predicted both closed and inactivated hERG conformations (*14, 15*). Here, the authors went beyond a “blind” AF2 prediction: they engineered structural templates to guide the network toward functionally relevant conformations, effectively steering it to sample gating-relevant structural states that pure homology or MSA-based prediction might miss. Their models exhibited plausible pore dimensions, key gating residue orientations, and features consistent with known mutational and electrophysiological effects, lending confidence that these *in silico* states may approximate true physiological conformations. Another AI-based modelling program, Chai-1, offers an alternative approach that can operate in single-sequence mode, drawing on large-scale protein language models to infer structural features without reliance on MSA (*16*). AF2 and Chai-1 are different AI models with different architectures and training data, meaning that for any given protein they might predict a slightly different structure. To the best of our knowledge, Chai-1 has not yet been applied to modelling any human ion channels, aside from our own recent work on modelling synthetic cannabinoid receptor agonists (SCRAs) (*17*) .

Given the importance of accurate closed-state models for pharmacological screening, we sought to independently validate and extend these recent AF2-based predictions using an alternative AI-driven framework. We generated a closed-state hERG model using Chai-1 in an ‘out-of-the-box’ manner (i.e. generating 5 models using the default settings), and compared it to the AF2-derived closed model, a homology model built for this study based on the *R. norvegicus* EAG channel (PDB: 5K7L), and two experimentally resolved structures, representing the open (PDB 9CHP) and non-conductive or inactivated state (PDB:9CHQ). We assessed these models using a combination of structural and dynamic metrics, including pore geometry, helix dynamics, water and ion permeation (via all-atom molecular dynamics), and lipid interactions (via coarse-grained molecular dynamics).

Our findings reveal a high degree of structural similarity between the Chai-1 and AF2 closed-state models, reinforcing the validity of both as plausible representations of the channel’s closed conformation. We also identify state-dependent differences, including helix dynamics and lipid recruitment. These results further highlight the growing potential of AI-based modelling to identify previously uncharacterised, pharmacologically relevant states of membrane proteins (*18*–*22*), thereby enhancing both mechanistic understanding and predictive drug-safety evaluation.

## Results

### Use of AI modelling to generate a closed model of the hERG channel

All five Chai-1 hERG models exhibit consistently high confidence, particularly within the core region of the channel. All models show strong inter-chain predicted TM-scores (ipTM ≈ 0.83), indicating reliable complex assembly, while the predicted local accuracy (pLDDT) is especially high throughout the complex, especially the pore region (**Figure** S1), supporting the structural integrity of the key functional domains. Structurally, the Chai-1 models closely resemble the previously published template-based AF2 model (*14*), with particularly strong agreement in the central pore architecture. Although some variability is observed in VSD domain and C2 linker, the pore domains remain highly conserved across models, with root-mean-square deviations (RMSDs) of approximately 0.53-0.57 Å (**Figure** S2). Notably, this agreement is better than that observed with the *R. norvegicus* cryo-EM EAG1 closed state structure (PDB: 5K7L (*8*)), which is commonly used (*9*) as a template for human hERG modelling, where RMSDs range from ∼0.63 to 0.86 Å (**Figure** S3). Taken together, the high ipTM and pLDDT scores, combined with the low RMSD relative to the AF2 model, support the robustness of the Chai-1 predictions. Based on these metrics, model 2 was selected for all subsequent analyses (**Figure** 2A).

**Figure 2:**
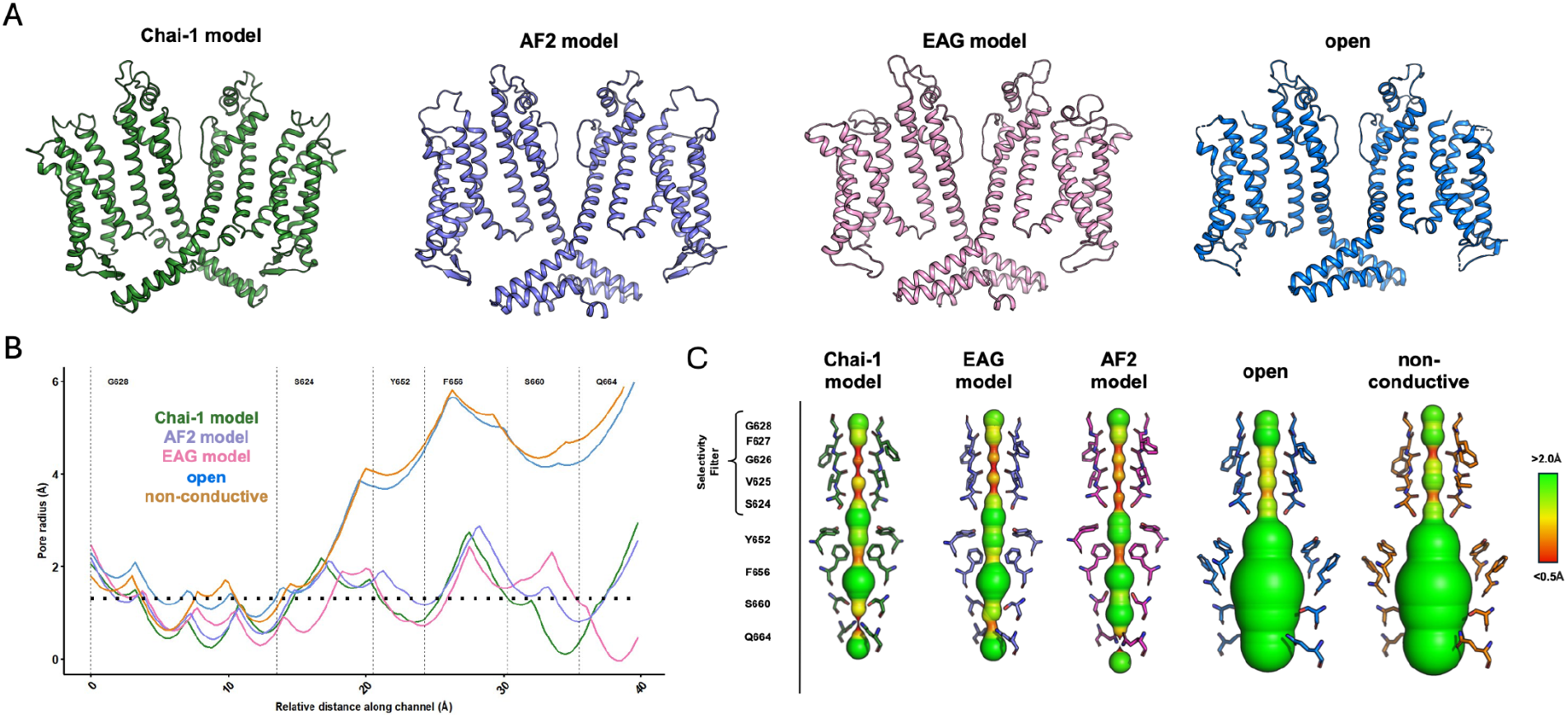
Comparison of Chai-1 models with previously reported hERG structures. (A) Structural alignment of Chai-1 model 2 with the previously published template-based AF2 model (“AF2 model”), a homology model based on the EAG structure (“EAG model”: based on PDB 5K7L), and the open-state cryo-EM structure (“open”: PDB 9CHP). (B) Pore radius profiles calculated using HOLE for the structures shown in panel A, with the addition of a non-conductive cryo-EM structure for comparison. (C) Close-up view of key pore-lining residues across all models, shown in the context of their corresponding HOLE profiles.

### Pore analysis of the Chai-1 closed model

The selected Chai-1 model exhibits a pore profile that is broadly consistent with a closed state across key gating regions (**Figure** 2B). In the selectivity filter region (around G628-S624), the pore radius in the Chai-1 model drops below ∼1.5 Å, approaching or falling beneath the approximate radius required for K^+^ permeation (indicated by the dashed line). This suggests that the filter adopts a constricted conformation that limits K^+^ passage. Comparison to the closed AF2 model and the closed EAG-based homology model suggests a very high degree of similarity in this region, including in the orientation of key selectivity filter residues (**Figure** 2C) supporting its designation as a closed state. The open and non-conductive cryo-EM structures are generally wider in this region, reaching above 1.5 Å .

Moving into the central cavity (around the canonical binding residues: Y652-F656), the Chai-1 model shows a moderate expansion, with the width increasing to ∼2.5-3 Å, again in line with the AF2 model and EAG-based homology model, and far narrower than the open and non-conductive cryo-EM structures, both of which display substantially wider cavities (>4 Å) (**Figure** 2B). The side chains of Y652 and F656 closely match those of the AF2 model and EAG homology model, with both pointing into the conduction pathway (**Figure** 2C). Notably, a constriction in the Chai-1 model occurs toward the intracellular gate region (around S660-Q664). Here, the pore radius sharply decreases to near 0-1 Å, representing a clear steric occlusion (**Figure** 2B). This is driven by the side chain position of Q664 (**Figure** 2C). This is substantially narrower than the AF2 model, and mimics a nearby constriction in the EAG model, also caused by Q664 (**Figure** 2C).

Overall, the Chai-1 pore profile appears to combine a constricted selectivity filter with a tightly closed intracellular gate. This dual constriction is characteristic of a closed state, and despite subtle differences, shows overall a high similarity to the AF2 and EAG homology models in the degree and location of pore narrowing.

### Analysis of the VSD in the Chai-1 model

Comparison of the voltage-sensing domain (VSD) across models shows that the overall architecture is well preserved, with strong agreement between the Chai-1 models and the previously reported AF2 structure (**Figure** 3A), and slightly lower similarity to the EAG-based homology model (**Figure** 3C). The most prominent structural deviation is localised at the C-terminal end of the S4 helix, immediately preceding the S4-S5 linker, a region known to play a critical role in coupling voltage sensing to pore opening.

**Figure 3:**
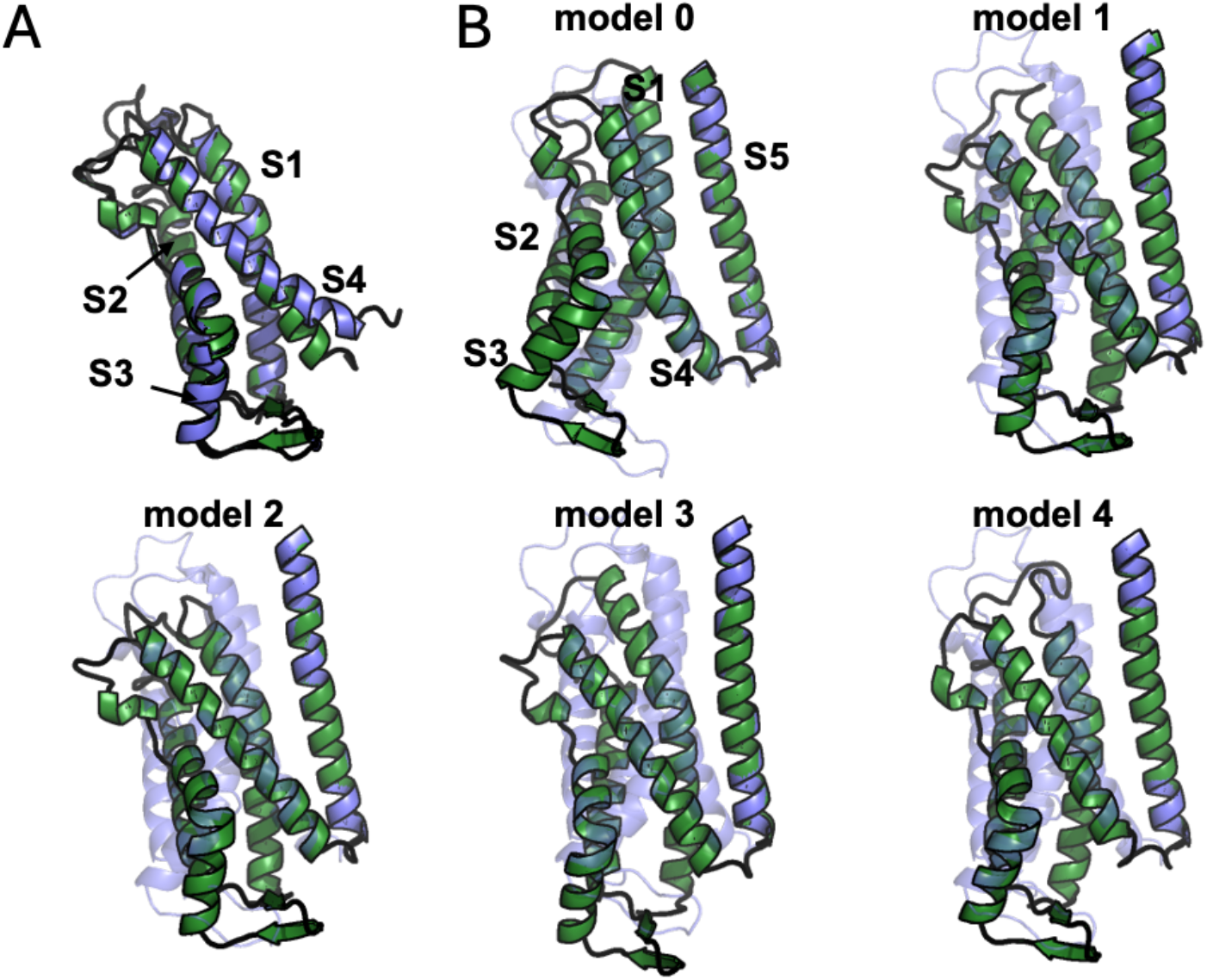
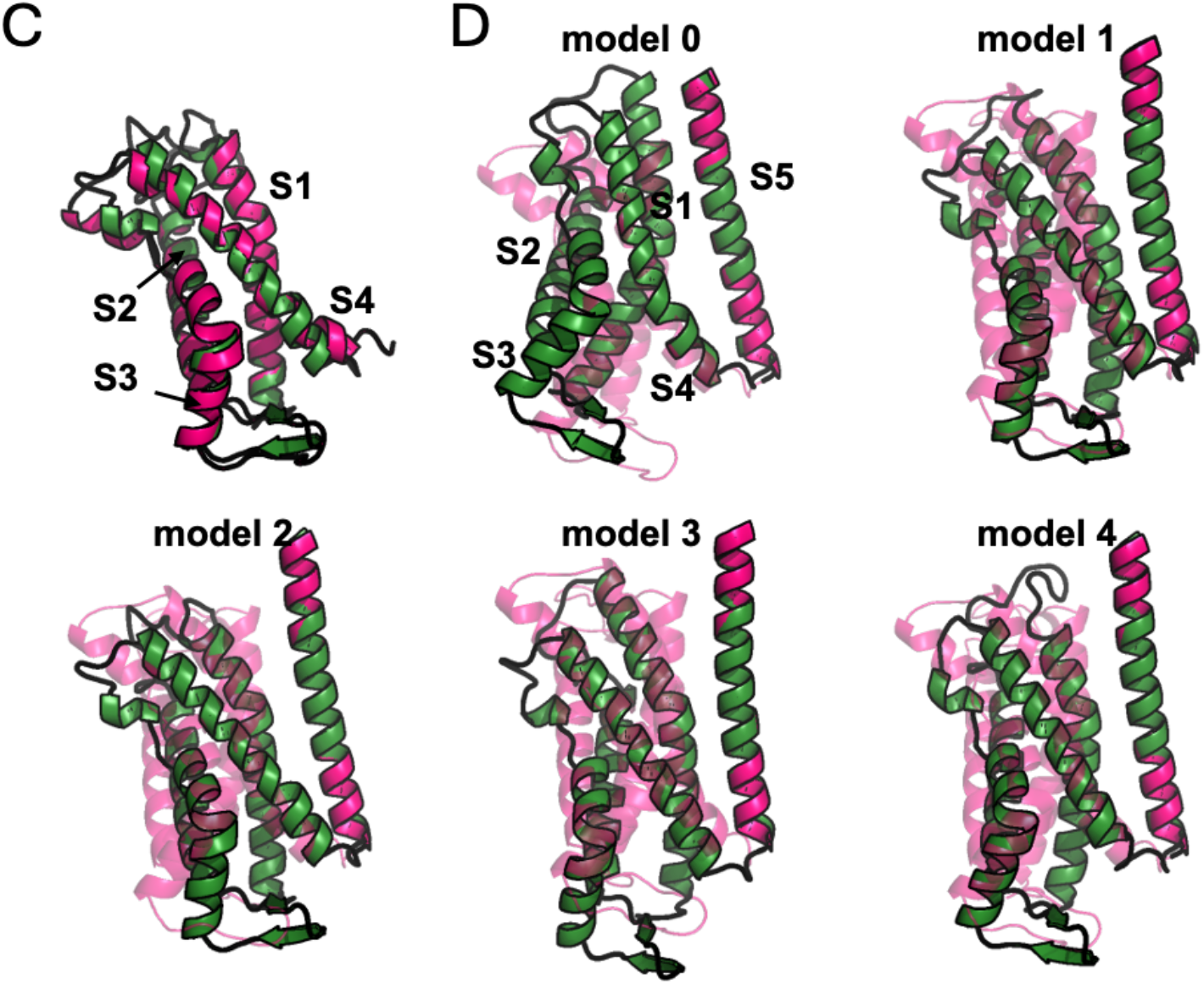
Comparison of the VSD and adjacent S5 helix between models. A) VSD of selected Chai-1 model (green) aligned to AF2 model VSD (blue), producing an RMSD of 1.1 Å. The other Chai-1 models show similar conservation (1.2, 1.0, 1.0, and 1.1 Å respectively). B) Views of all five Chai-1 VSD domains and adjacent S5 (green) compared to the AF2 model (transparent blue), as aligned on S5. C) As panel A, but for the Chai-1 model and the EAG homology model (pink). RMSD = 1.5 Å, and 1.6, 1.5, 1.5 and 1.4 Å for the other Chai-1 models D) As panel A, but for the EAG homology model (transparent pink).

Consistent with this, the relative orientation of the VSD with respect to S5 varies substantially between models. All five Chai-1 models display differences in this interface, indicating a degree of conformational heterogeneity. Notably, Chai-1 model 0 closely recreates the AF2-derived orientation, whereas models 1-4 adopt a markedly different configuration, characterised by a pronounced downward tilt of the VSD relative to the pore domain (**Figure** 3B). This shift likely reflects alternative positioning of the S4-S5 linker and may correspond to distinct coupling states between the VSD and pore. In contrast, none of the Chai-1 models reproduce the VSD orientation observed in the EAG-based homology model (**Figure** 3D). Collectively, these observations reinforce the idea that, while the core VSD fold is conserved, its orientation relative to the pore domain is highly variable, with likely functional impacts on channel gating.

### Stability and dynamics of the hERG Models

We next performed atomistic MD simulations to assess the stability and dynamics of the Chai-1 model, comparing it against the AF2 model, the EAG-based homology model, and the open cryo-EM structure. For each system, we ran five independent 1 µs replicates in a fully solvated lipid membrane. Root-mean-square deviation (RMSD) analysis of the protein backbone revealed that while all hERG models were generally stable, the two experimental structures exhibited greater overall stability than the Chai-1 and AF2 predictions, especially for 1 or 2 replicates of each which showed higher dynamics (>0.6 nm RMSD) (**Figure** S4).

Root-mean-square fluctuation (RMSF) analysis further indicated that the core motifs of the hERG channel remained generally stable across all simulations (**Figure** 4A). Notably, the S2-S3 linker in the open state was more dynamic than in the closed states, whereas all three closed states exhibited higher flexibility in the S3 helices compared to the open state. The EAG-based model showed a significantly more dynamic S4 helix than all other states, while the open state displayed a more dynamic S5. The turret helix was consistently dynamic across all states, however, the pore helix was markedly stabilised in the open state relative to the closed states. In all systems, the C-linker (C2) was the most dynamic region (**Figure** 4A-B), presumably contributing to the higher RMSD in some simulations. Overall, these data reveal distinct conformational flexibility profiles between the closed and open states of the hERG helices

**Figure 4:**
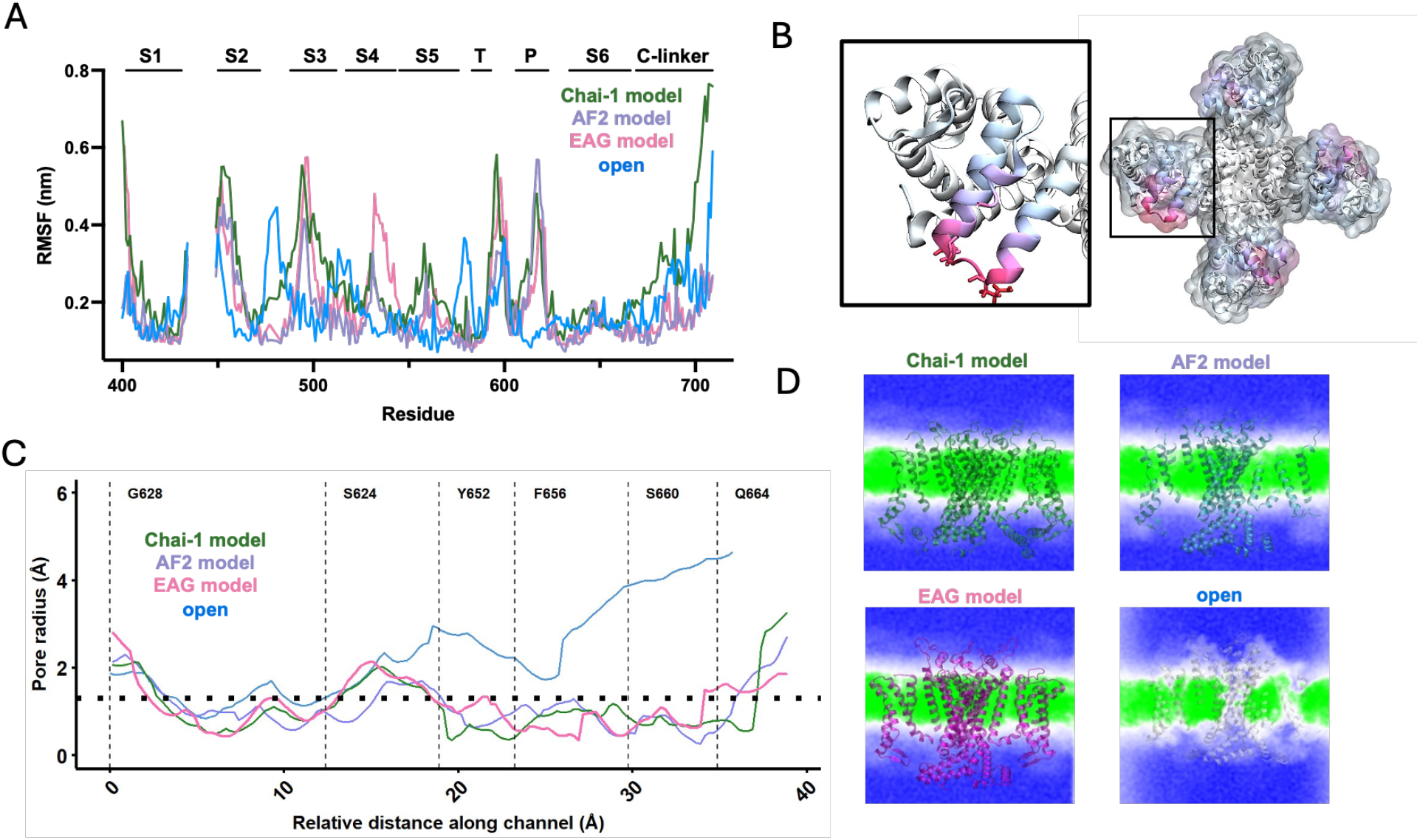
Molecular dynamics of the hERG models. (A) Per-residue RMSF analysis along hERG, averaged across four subunits for 5 x 1 µs of MD simulation. S1-6 helices are labelled, along with the turret (T) and pore (P) helices and C-linker. B) RMSF values projected onto the Chai-1 model, with closeup of the C2 linker inset showing high flexibility in this region. C) HOLE traces as run on post simulation snapshots for each of the hERG models D) Densities of waters (blue) and lipids (green) computed over 5 × 1 µs of MD simulation for each model using VMD.

### Analysis of the hERG pore during MD

HOLE analysis of post-MD snapshots (**Figure** 4C) revealed that the pore dimensions remained relatively consistent with the initial input models (**Figure** 2B), though subtle refinements were observed. Firstly, the narrow constrictions in the Chai-1 and EAG models around the ca. 35-40 Å mark are now less pronounced. Further, the widening in all of the input closed models at the ca. 28 Å (between F656 and S660) reduces from about 3 Å to less than ∼1.5 Å. Finally, the open state narrows considerably, especially around F656, from about 6 Å to 2-3 Å. Together, this shows that all input models, including the experimentally-derived open cryo-EM structure, are subject to change during MD simulations, highlighting the utility of complementary simulation methods when probing structural models.

During the simulations, water molecules moved freely within and around the hERG models. To characterize the hydration state of the channels, we calculated water molecule densities across all MD frames (**Figure** 4D). As expected, no water density was observed within the pore for the three closed-state models (Chai-1, AF2, and EAG-based homology model), whereas substantial water flux was maintained in the open model. This confirms that all predicted closed models remain in non-open, dehydrated state throughout the 5 x 1 µs MD simulation.

### Lipid binding sites in the different hERG states

A previous study (*23*) used MD to probe lipid interactions with the hERG channel using an open hERG model (*7*). We reasoned that it would be useful to re-evaluate these interactions using the two recent closed-state predictions (our Chai-1 and the previously published AF2) and the newer open-state cryo-EM structure (PDB 9CHP (*6*)). Furthermore, whereas previous work employed the Martini 2.2 force field (*24*), we utilised the updated Martini 3.0 (*25*).

All models were subjected to five independent 10 μs coarse-grained MD simulations in a mixed lipid bilayer containing 20% ceramide (Martini lipid “DPCE”) and 5% of the polyunsaturated lipid (C18:2) dilinoleoyl phosphatidylcholine (DIPC; Martini “DIPC”). Previous work identified M651 as a key residue involved in ceramide binding, so our initial analysis focused on this site. Across simulations, we observed substantial enrichment of ceramide in the vicinity of the channel, as evidenced by both lipid occupancy analysis using the PyLipID package (*26*) (**Figure** 5A) and spatial density maps calculated in VMD (**Figure** S5).

**Figure 5:**
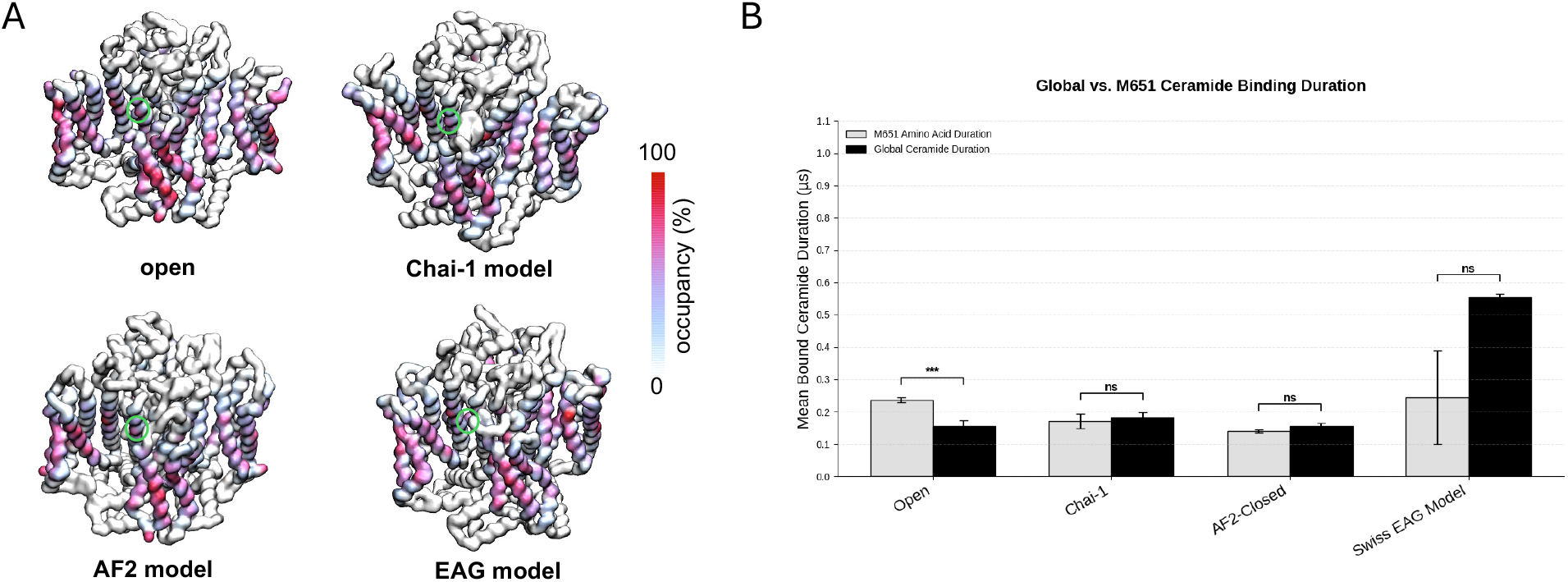
Coarse-Grained MD and lipid Interactions. (A) PyLipID-derived occupancies for ceramide, projected on the hERG structures. M651 position is denoted with a green circle. (B) Quantification of binding duration between ceramide and residue M651 on each of the hERG channel models in relation to general ceramide binding duration. Global values are taken as an average duration from all sites identified with PyLipID.

Kinetic analysis of lipid interactions using PyLipID revealed a significant enrichment of ceramide around M651 in the open-state model (**Figure** 5B), consistent with the prior findings (*23*). In contrast, no significant enrichment was detected at this site in any of the closed-state models, including our Chai-1 model, indicating a clear state dependence of this interaction (**Figure** 5B). No significant enrichment was observed for the polyunsaturated DIPC lipid in any model (**Figure** S6). Together, these results reinforce the idea that ceramide binding at M651 is both selective and conformation-dependent, with potential implications for lipid-mediated modulation of hERG gating.

## Discussion

Ion channels, such as hERG, do not exist in a single static conformation but instead occupy a continuum of interconverting microstates – indeed, this underpins their complex gating behaviour. As such, no single structure can fully capture the functional landscape of the channel, and the availability of multiple structurally and dynamically distinct models provides a more comprehensive framework for understanding ion channel function. In this area, AI-based modelling tools can be of considerable value. AF2 has previously been previously used to explore NaV channel conformational states, finding that the output models represent multiple channel conformations, including both experimentally-observed and novel conformations (*13*). Another study comparing AF2 with RosetTTAFold (*27*) and ESMfold (*28*) suggest these methods capture the voltage-sensing and pore domains of voltage-gated ion channels with high confidence and in a number of states, although they highlight uncertainties of modelling unstructured loop and termini regions (*12*). Finally, a template-based AF2 modelling study of hERG demonstrated their approach was able to predict a range of experimentally unresolved hERG states, providing insights into hERG inactivation mechanisms and uncovering molecular features relating to drug binding (*14*).

Our results highlight the utility of a different AI-based modelling package, Chai-1 (*29*), as another powerful tool for predicting functionally relevant conformational states of ion channels, particularly in cases where such states are not readily captured by default structure prediction pipelines. Our Chai-1 derived hERG model adopts a closed pore architecture, as evidenced by the pronounced constriction at the intracellular gate and the overall HOLE profile. This is notable because standard AF2 predictions typically favour more open or intermediate conformations, and obtaining a fully closed hERG structure typically requires extensive intervention, including template biasing, multiple sequence curation, or iterative refinement workflows. In contrast, Chai-1 appears capable of natively sampling this important closed state without the need for such highly involved pipelines. This commends the utility of AI-based modelling, where previously modelling of the hERG closed state required extensive and complex modelling pipelines (*30, 31*).

Importantly, while the Chai-1 model shows strong overall agreement with both the AF2-derived and EAG-based homology models, the subtle but consistent differences in pore geometry – particularly within the central cavity and intracellular gate – may reflect a distinct conformational state. Given the well-established state dependence of hERG drug binding, these variations could have meaningful implications for drug interactions, especially for compounds that preferentially bind to the closed state. As such, our Chai-1 structural model expands the available structural repertoire for hERG, offering an additional and potentially more physiologically relevant template for future structure-based drug design and *in silico* screening efforts.

### Structural changes within the pore and VSD

From a mechanistic perspective, differences observed within the voltage-sensing domain (VSD) are of particular interest in the context of channel activation. AI structural modelling represents a powerful method for generating insights into the dynamics of this region, as demonstrated in a recent study using AF2 to produce a large number of VSD structural states, including activated, deactivated, and intermediate conformations (*32*).

Our Chai-1 modelling recapitulates the overall architecture of the VSD observed in both experimental structures and AF2-based models; however, the relative orientation of the domain shows notable variation (**Figure** 3). The data indicate that, while the global VSD fold is conserved, shifts in orientation may reflect alternative conformational states or coupling relationships between voltage sensing and pore gating. In particular, we observe substantial movement of S4 relative to S5, driven by rearrangements in the S4-S5 linker. Given that the S4 helix serves as the primary voltage sensor, undergoing outward and rotational motion in response to membrane depolarisation (*33*), such variability is likely to have direct implications for channel activation. The S4-S5 linker is well established as a key mechanical transducer that couples VSD motion to opening of the intracellular gate (*34*), and subtle differences in its conformation may therefore bias the channel toward distinct gating states. In this regard, the structural plasticity of this region in our Chai-1 models, along with the increased flexibility in the MD in the EAG-based model (**Figure** 4) may reflect functionally relevant heterogeneity within this coupling interface. However, we note that the pLDDT of these regions is lower than the central pore, raising the possibility that some of the observed variability arises from reduced model confidence. This highlights a broader challenge in interpreting AI-derived structures, namely distinguishing genuine conformational diversity from uncertainty in poorly constrained regions.

Additional differences in dynamics are observed in MD simulations within regions implicated in VSD-pore coupling. The increased mobility of the S2-S3 linker in the open-state model is not straightforward to interpret mechanistically but may influence the energetic landscape of S4 motion, potentially modulating how readily the voltage sensor transitions between states and how effectively these movements are transmitted to the pore domain. This linker has also been proposed to be important for proper transduction of VSD rearrangements to opening and closing the cytoplasmic gate (*35*), so increased flexibility here could reflect a loosening of these interactions in the open state. Conversely, the reduced flexibility of the pore helix in the open state suggests a stabilising role once the conductive conformation is achieved. A more rigid pore helix may help maintain precise coordination of permeating ions and preserve the geometry of the selectivity filter, both of which are essential for efficient conduction. In contrast, increased flexibility in closed states may reflect a less constrained environment in the absence of ion flux.

### State-dependent lipid binding at the hERG channel

Miranda et al. (*23*) previously investigated the regulation of channel gating by ceramide species, demonstrating that specific sphingolipids can directly modulate hERG function. Using coarse-grained simulations, their analysis focused primarily on the open-state conformation of hERG and identified putative lipid interaction sites centred around M651. In the present study, we revisited this question using an updated coarse-grained force field (*25*), alongside a more recent open-state hERG model and our newly generated closed-state Chai-1 model. Our data successfully reproduced their key findings, and further revealed that ceramide enrichment at the M651 site is a feature of the open state and is largely absent in the closed conformation (**Figure** 5). This suggests that conformational changes associated with channel opening either expose or stabilise a ceramide-binding site that is not accessible in the closed state, supporting the specificity of ceramide interactions in regulating hERG gating.

Notably, we have previously identified this same region as a binding site for SCRAs (*17*) and the pollutant phenanthrene (*36*), where ligand binding at this site leads to channel block. Together, these observations point to a possible mechanistic link between lipid binding at M651 and modulation of pore gating, with implications for both endogenous regulation and drug-induced hERG inhibition.

### Limitations

While modelling metrics such as pLDDT and pTM scores can be used to assess model confidence, the use of MD provides a more robust cross state and cross method comparison (*37*). Here, we used molecular dynamics (MD) simulations to both test our AI-generated model and probe its dynamic behaviour, including protein dynamics, pore structure, water permeation, and lipid interactions. Our data support previous work suggesting that combining AI modelling with MD is a powerful combination for interrogating ion channel structure and function (*14, 38, 39*). We note that experimental analysis is also of utmost importance for validating newly predicted states, aiming to design experiments that discriminate between these states and known structures (*40*).

However, several limitations should be considered. AI-based protein structure prediction methods, including Chai-1 and AF2, are inherently constrained by their training data and may not fully capture the full conformational landscape of highly dynamic systems such as ion channels. Similarly, MD simulations are limited by accessible timescales and may not adequately sample rare but functionally important transitions. As such, the structures and dynamics presented here should be interpreted as representative snapshots within a broader ensemble. Within this context, the Chai-1 model and associated MD data should be viewed as a valuable addition to the existing toolkit, complementing experimental structures and other computational models in probing the structural and pharmacological properties of hERG.

### Future and clinical outlook

Looking forward, the integration of Chai-1 and related models into drug discovery pipelines offers significant potential. The availability of additional hERG structural states, including the closed conformation described here, expands the toolkit for structure-based screening and may improve early prediction of cardiotoxic liabilities. Given the central role of hERG dysfunction in drug-induced arrhythmias, particularly Long QT syndrome, such advances have the potential to enhance the safety of therapeutic development and, in the longer term, contribute to better outcomes for patients at risk of life-threatening cardiac events.

The future release of a closed-state hERG cryo-EM structure would provide an important benchmark; however, such a structure would still represent only a single conformational snapshot. In this regard, AI-derived models, particularly when coupled with MD simulations, are well positioned to capture aspects of the dynamic behaviour that underpins the distinctive gating kinetics of hERG. Continued development and integration of these approaches may ultimately provide deeper mechanistic insight and improve our ability to rationally target this clinically important channel.

## Data availability

Chai-1 models and confidence metrics, along with post-MD snapshots, are available for download from the OSF (https://osf.io/gbejy/).

## Acknowledgements

This project made use of time on ARCHER2 granted via the UK High-End Computing Consortium for Biomolecular Simulation, HECBioSim (http://www.hecbiosim.ac.uk), supported by EPSRC (grant no. EP/R029407/1). This work was carried out using the computational facilities of the Advanced Computing Research Centre, University of Bristol - http://www.bristol.ac.uk/acrc/. CU is supported by a BHF PhD Programme. TO is supported by a BHF Non-clinical PhD Studentship (FS/PhD/25/29771). RC and JH are supported by the British Heart Foundation (FS/PhD/25/29771).

## Methods

### Production of hERG models

To model the hERG channel, we deployed the novel protein-folding software Chai-1(*16*) using the webserver (https://lab.chaidiscovery.com/). To model the core channel, we used a truncated amino acid sequence of hERG, containing the trans-membrane domain (residues 400-718). We used the default settings, with now MSAs and restraints applied. In parallel, we used SwissModel (https://swissmodel.expasy.org/) (*41*) to produce a tetrameric model of hERG residues 401-718 hERG using the rat EAG closed structure (PDB 5K7L) (*8*). Alongside these, we obtained the recently published closed AF2 structure (trimmed to residues 398-719) (*14*) and the experimentally determined open hERG structure (PDB:9CHP; residues 398-709) (*6*), Visual analysis of the channels was performed using PyMOL (Schrödinger LLC).

### Analysis of models

To help characterise the conformational state of the Chai-1 predicted hERG structures and post-MD snapshots (1 µs), the programme HOLE was used to measure pore radius, using a Monte Carlo simulated annealing procedure to detect an optimum route through a channel (*42*). Visual analysis of the HOLE profiles was performed using PyMOL (Schrödinger LLC).

### Atomistic MD simulations

MD simulations were conducted to assess the stability of our hERG models. Proteins were described using the CHARMM36m forcefield (*43*). Each hERG model was placed into a lipid bilayer consisting 70% 1-palmitoyl-2-oleoyl-sn-glycero-3-phosphocholine (POPC), 20% cholesterol, and 10% palmitoylsphingomyelin (PSM). Membranes were solubilised with TIP3P water and neutralised with potassium chloride ions at concentration 0.15M via CHARMM-GUI Membrane Builder (*44, 45*). Protein side chains were all set to default protonation states.

The systems were minimised and equilibrated using the standard CHARMM-GUI protocol, with an additional 20 ns of NPT equilibration with xyz positional restraints of 50 kJ/mol/nm^2^ applied to the protein backbone. MD simulations were then conducted across each system for 1 µs with a total of 5 repeats. Simulations were run using GROMACS 2024 (*46, 47*). Simulations used a 2 fs time step in the NPT ensemble with the velocity-rescale thermostat and c-rescale pressure coupling. The results of the MD simulations were analysed using built-in GROMACS tools. The results of the MD simulations were analysed using built-in GROMACS tools and in-house scripts. Visual analysis was performed using PyMOL (Schrödinger LLC) and VMD (*48*).

### Coarse-grained MD simulations

Coarse-grained molecular dynamics simulations were built for each of the hERG models. Protein atoms were converted to the CG Martini 3 force field (*25*) using the martinize method (*49*). Additional bonds of 500 kJ mol^−1^ nm^−2^ were applied between all protein backbone beads within 0.9 nm. Proteins were built into membranes composed of 40% POPC (palmitoyl oleyl phosphatidylcholine), 10% POPE (palmitoyl oleyl phosphatidylethanolamine), 20% DPCE (ceramide), 20% cholesterol, 5% DIPC (dilinoleoyl phosphatidylcholine), and 5% POPS (palmitoyl oleyl phosphatidylserine), using the insane method (*50*). All systems were solvated with Martini waters and Na^+^ and Cl^−^ ions to a neutral charge and 0.15 M. Systems were minimized using the steepest descents method, followed by 1 ns equilibration with 5 fs time steps, then by 100 ns equilibration with 20 fs time steps, before 5 × 15 µs production simulations using 20 fs time steps, all in the NPT ensemble at 323 K with the V-rescale thermostat (τ=1.0 ps) and semi-isotropic Parrinello-Rahman pressure coupling at 1 bar (τ=12.0 ps). The reaction-field method was used to model long-range electrostatic interactions. Bond lengths were constrained to the equilibrium values using the LINCS algorithm. Density analysis was performed using the VolMap tool of VMD, with the default settings (*48*). Lipid binding sites and lipid-residue interactions were determined using the PyLipID package, which provides both occupancy and residence time data (*26*). Reported occupancy and residence time values are taken analysis of the total simulation time. Simulations were run using GROMACS 2024 (*46, 47*).

