## Supplementary figures for "Integrating AI and molecular modeling for structural prediction of a closed state of the hERG channel"

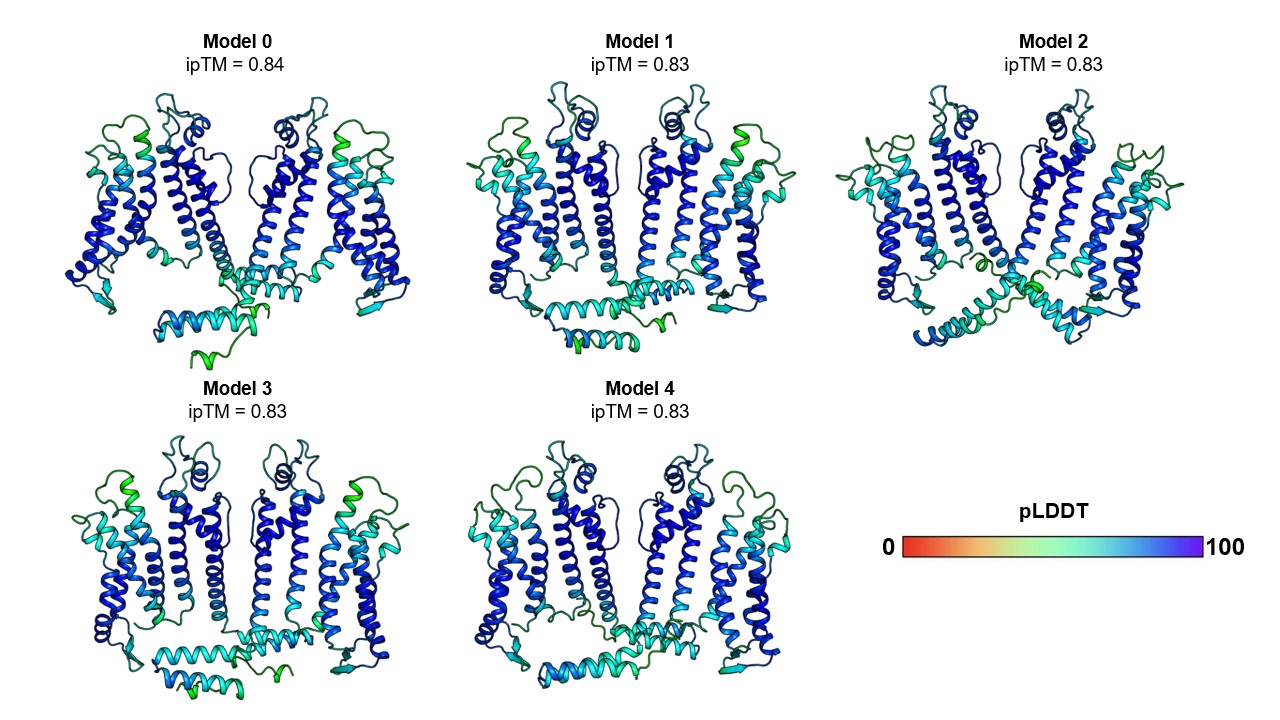

**Figure S1: Chai-1 pLDDT for all 5 models**

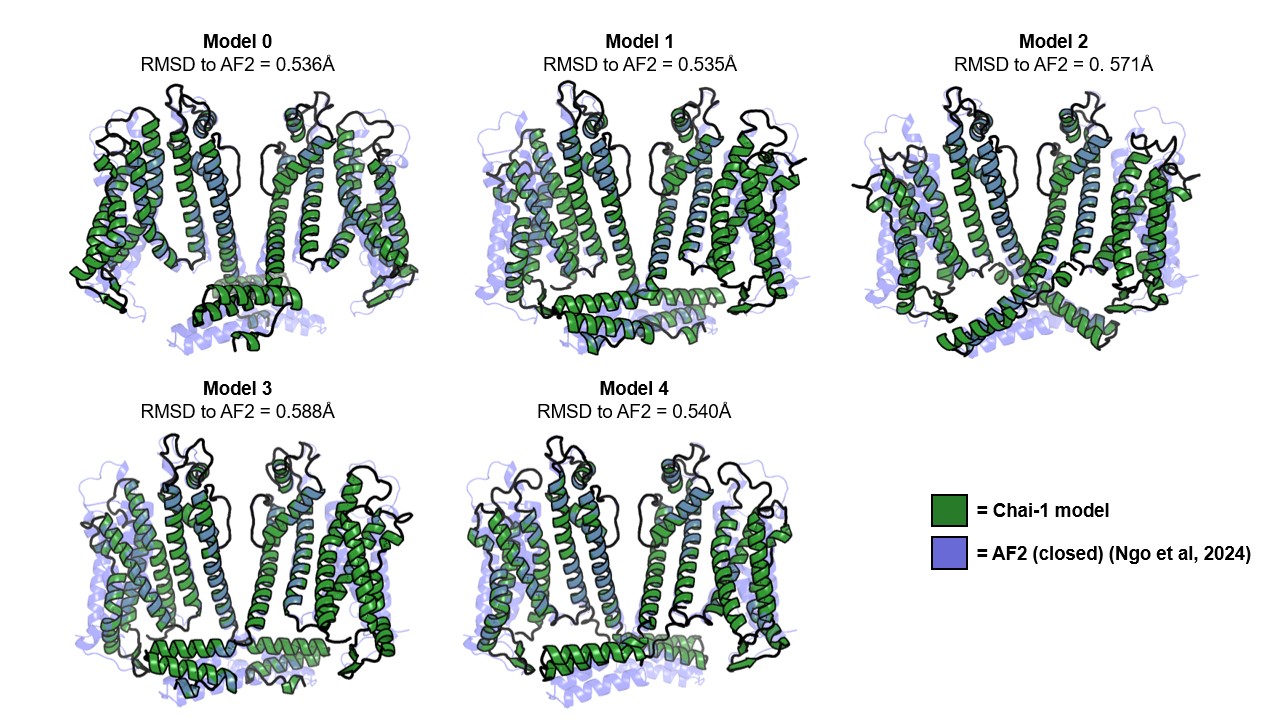

**Figure S2: Alignment of all 5 Chai-1 models to the previously published AF2 closed structure**

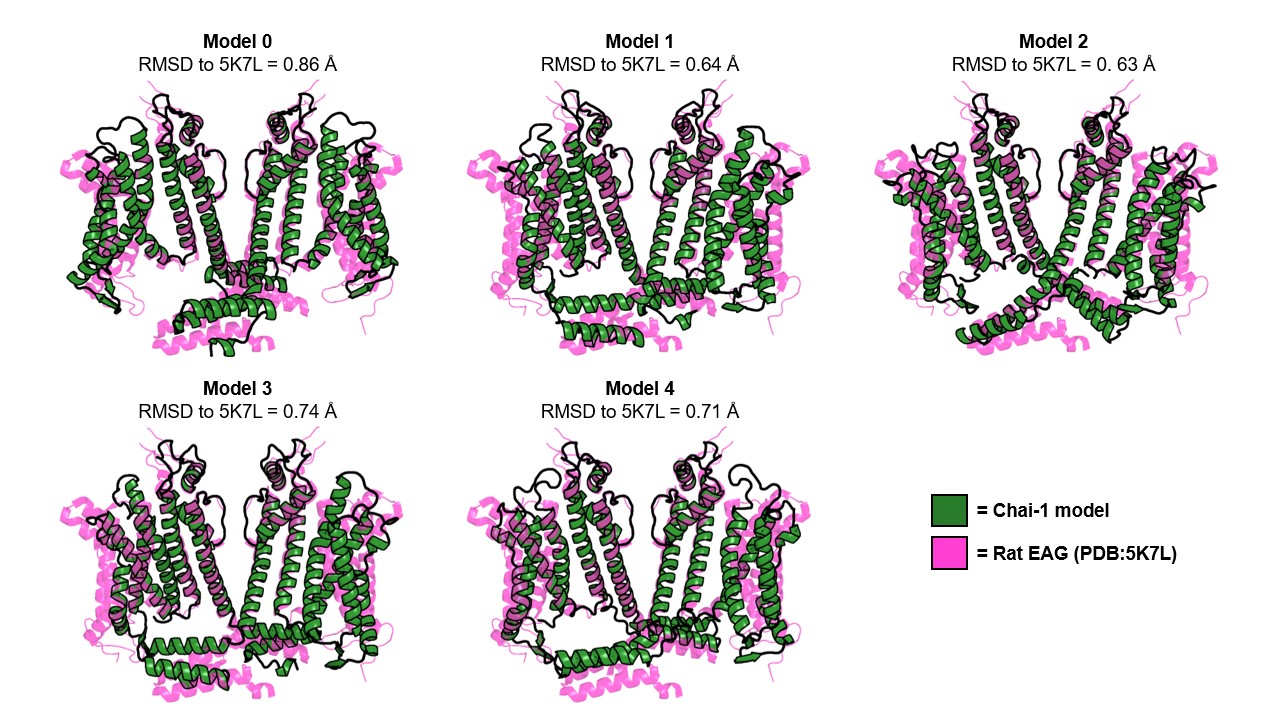

**Figure S3: Alignment of all 5 Chai-1 models to the previously published *R. norvegicus* EAG structure**

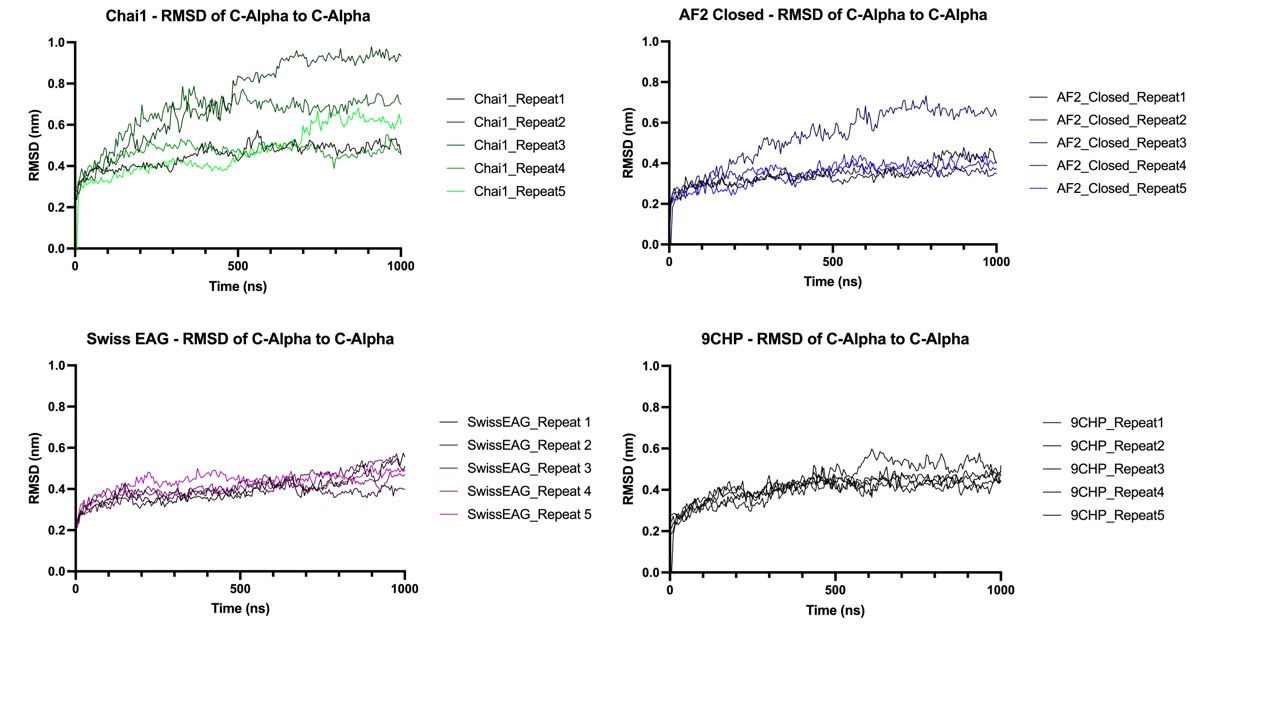
**Figure S4: Backbone RMSDs for the entire hERG channel across the atomistic MD simulations for each of the hERG models**

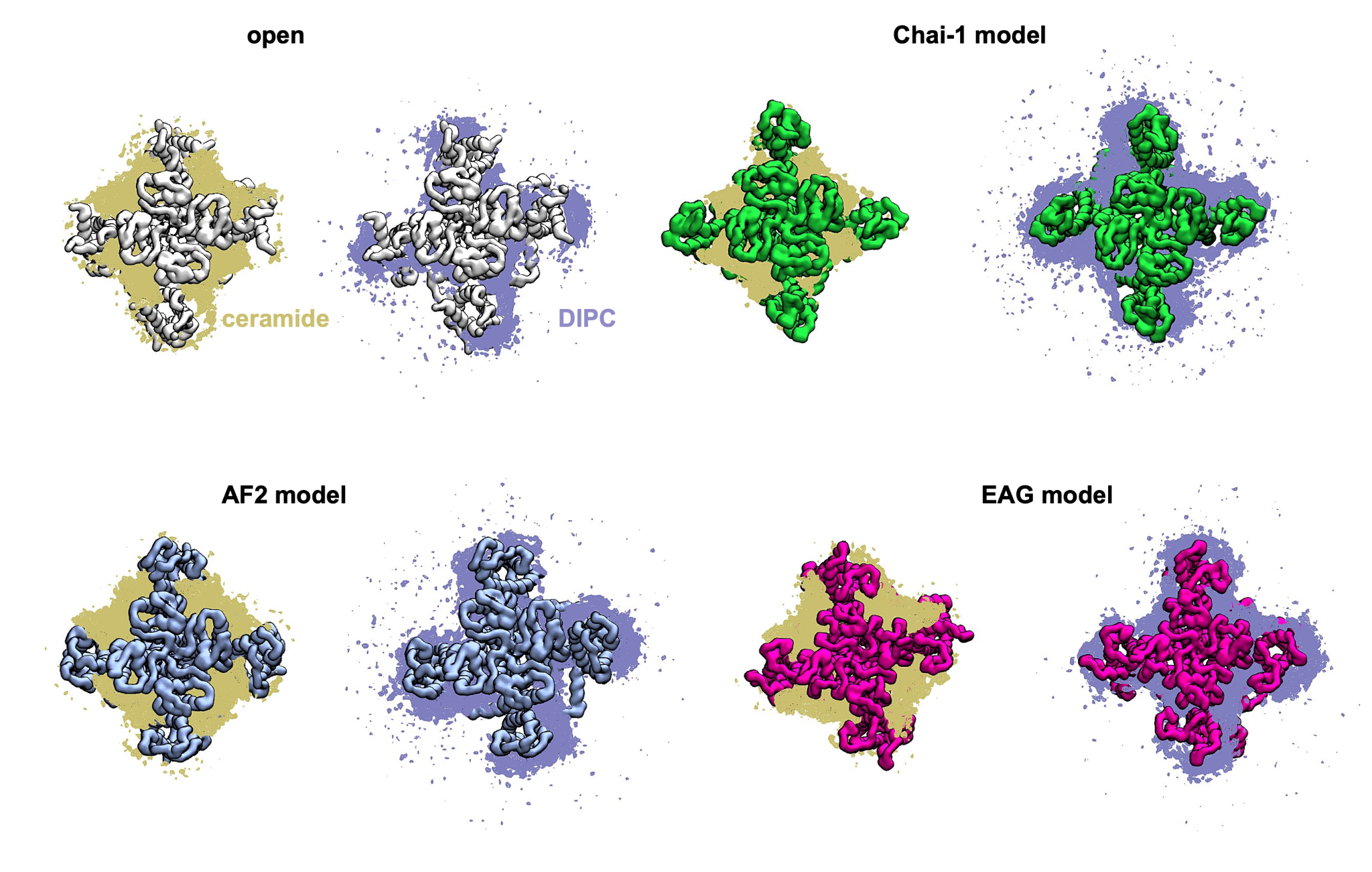

**Figure S5: Computed densities of ceramide in relation to each of the hERG models based on 5 x 10 µs of coarse-grained MD.** Densities computed using VMD VolMap. Gold relates to ceramide (DPCE) density. Blue relates to DIPC (dilinoleoyl PC) density.

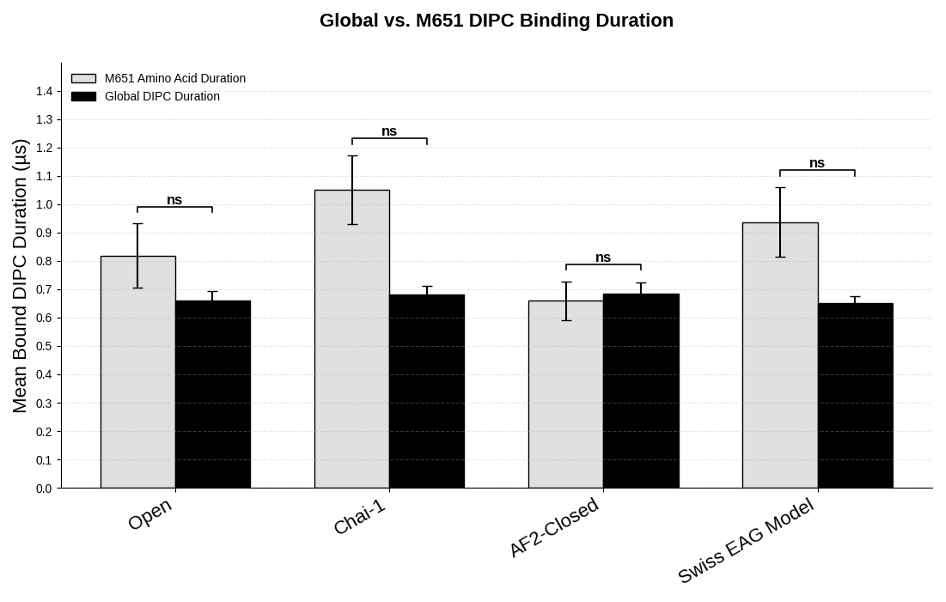

**Figure S6: Graph showing enrichment of DIPC lipid at residue M651 during CG-MD of the various hERG models.**
